# Immunity to seasonal coronavirus spike proteins does not protect from SARS-CoV-2 challenge in a mouse model but has no detrimental effect on protection mediated by COVID-19 mRNA vaccination

**DOI:** 10.1101/2022.10.25.513804

**Authors:** Fatima Amanat, Jordan Clark, Juan Manuel Carreño, Shirin Strohmeier, Philip Meade, Disha Bhavsar, Hiromi Muramatsu, Weina Sun, Lynda Coughlan, Norbert Pardi, Florian Krammer

## Abstract

Seasonal coronaviruses have been circulating widely in the human population for many years. With increasing age, humans are more likely to have been exposed to these viruses and to have developed immunity against them. It has been hypothesized that this immunity to seasonal coronaviruses may provide partial protection against infection with severe acute respiratory syndrome coronavirus 2 (SARS-CoV-2) and it has also been shown that coronavirus disease 2019 (COVID-19) vaccination induces a back-boosting effects against the spike proteins of seasonal betacoronaviruses. In this study, we tested if immunity to the seasonal coronavirus spikes from OC43, HKU1, 229E or NL63 would confer protection against SARS-CoV-2 challenge in a mouse model, and whether pre-existing immunity against these spikes would weaken the protection afforded by mRNA COVID-19 vaccination. We found that mice vaccinated with the seasonal coronavirus spike proteins had no increased protection as compared to the negative controls. While a negligible back-boosting effect against betacoronavirus spike proteins was observed after SARS-CoV-2 infection, there was no negative original antigenic sin-like effect on the immune response and protection induced by SARS-CoV-2 mRNA vaccination in animals with pre-existing immunity to seasonal coronavirus spike proteins.

**Importance:** The impact that immunity against seasonal coronaviruses has on both susceptibility to SARS-CoV-2 infection as well as on COVID-19 vaccination is unclear. This study provides insights into both questions in a mouse model of SARS-CoV-2.

## Introduction

Severe acute respiratory syndrome coronavirus 2 (SARS-CoV-2), a beta-coronavirus (β-CoV), first emerged in late 2019 and has since then caused the coronavirus disease 2019 (COVID-19) pandemic (1). SARS-CoV-2 is the fifth CoV to cause large scale outbreaks in humans. The four seasonal CoVs – α-CoVs 229E (2) and NL63 (Netherlands 63 (3, 4)) and β-CoVs OC43 (organ culture 43) and HKU1 (Hong Kong University 1 (5)) - have likely been circulating in humans for decades, potentially even longer, and cause a sizable percentage of common colds every winter (6-8). In addition, OC43 is suspected to have jumped from cattle into humans in the late 19^th^ century and to be the causative agent of the 1889/1890 ‘Russian flu’ pandemic (9). Infections with seasonal CoVs happen frequently and it is assumed that a large proportion of the human population, maybe even the entire population, has immune memory against these viruses derived from previous infections. While immunity to seasonal CoVs may wane over time leading to reinfection, a large proportion of the population is likely - at least partially - protected (10).

In has been hypothesized that this pre-existing immunity to CoVs, including antibodies to the spike protein, may provide a degree of protection against SARS-CoV-2 infection and COVID-19, especially in children (11). Rare individuals in fact have detectable cross-reactive anti-SARS-CoV-2 spike antibodies before exposure to SARS-CoV-2 (12). It has also been shown that cross-reactive antibodies are induced in individuals infected with SARS-CoV-2 or immunized with COVID-19 vaccines in a ‘back-boost’ or ‘original antigenic sin’-like manner (13, 14). Here, we are investigated the protective effect of immunity against the different seasonal CoV spike proteins against SARS-CoV-2 infection in a mouse model and we also explored if pre-existing immunity to the seasonal CoV spike proteins has a detrimental impact on the protective effect of COVID-19 vaccination.

## Results

### Seasonal CoV spike proteins induce strong specific immune responses in mice with detectable cross-reactivity within α-CoVs and β-CoV

The spike proteins of 229E, NL63, OC43, HKU1 and SARS-CoV-2 were expressed as soluble trimers and mice were vaccinated with these proteins with an oil-in-water emulsion adjuvant in a prime-boost regimen (**Figure 1A**). Control mice were vaccinated in the same manner but with an irrelevant recombinant antigen, recombinant influenza virus hemagglutinin (HA). Four weeks post-boost animals were bled and their immune response was assessed by enzyme-linked immunosorbent assay (ELISA). Sera from 229E vaccinated mice reacted strongly with 229E spike and reduced reactivity was also detected against the NL63 spike (**Figure 1B**). No reactivity was detected to any of the three β-CoV spikes. While NL63 spike seemed to be slightly less immunogenic, it produced similar results with strong reactivity to the homologous spike, reduced activity to the 229E spike and some, albeit reduced reactivity to the β-CoV spikes (**Figure 1C**). Vaccination with OC43 led to strong anti-OC43 spike reactivity, reduced reactivity to the HKU-1 spike followed by reactivity to the SARS-CoV-2 spike (**Figure 1D**).

**Figure 1:**
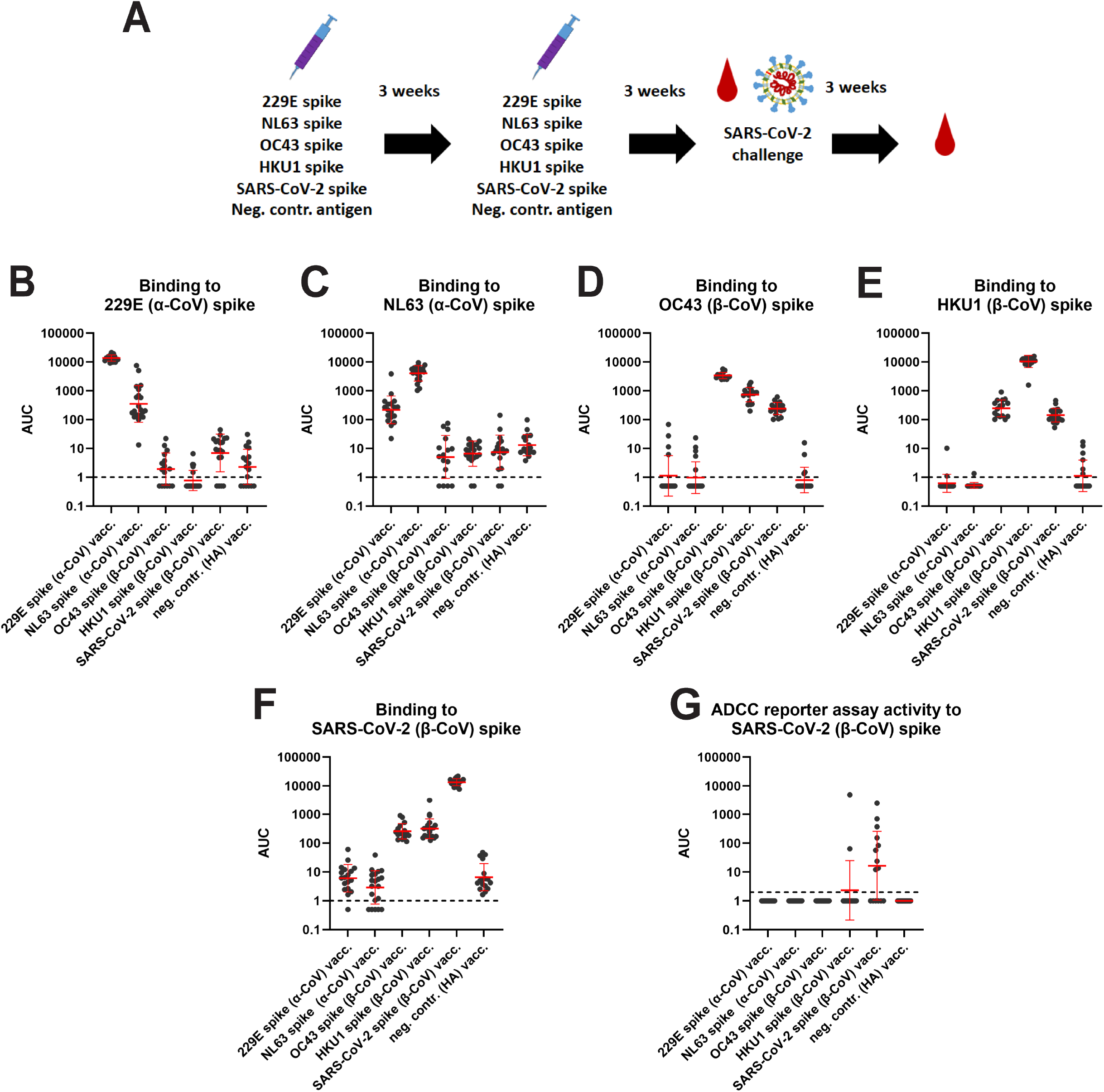
Antibody reactivity against CoV spikes after vaccination. **A** Schematic of vaccination regimen, challenge and sampling. **B** Serum reactivity to 229E spike protein. **C** Serum reactivity to NL63 spike. **D** Serum reactivity to OC43 spike. **E** Serum reactivity to HKU1 spike. **F** Serum reactivity to SARS-CoV-2 spike. **G** ADCC reporter activity to SARS-CoV-2 spike. 229E n=18, NL63 n=20, OC43 n-16, HKU1 n=20, SARS-CoV-2 n=19, neg. contr. n=17 – except for G where n=15 for all groups. Significant statistical differences are summarized in Supplemental Table 1. Red bars indicate geometric mean, the error indicates the geometric standard deviation. Samples were analyzed once except for G were the mean of two replicates is shown.

Reactivity to the α-CoVs was negligible. HKU1 spike vaccination induced strong reactivity to the homologous spike and lower titers to OC43 and SARS-CoV-2 spikes but reactivity to the α-CoV spikes was also at baseline (**Figure 1E**). Finally, vaccination with SARS-CoV-2 spike induced high titers to itself and some cross-reactivity to OC43 and HKU-1 but no reactivity to α-CoV spikes (**Figure 1F**). In addition to ELISA binding, we also assessed the activity of these sera in an antibody-dependent cellular cytotoxicity (ADCC) reporter assays. The results, shown in **Figure 1G**, indicate that a proportion of SARS-CoV-2 spike vaccinated mice in fact show activity in this assay against cell surface expressed spike protein. However, despite binding activity, sera from mice vaccinated with any of the seasonal CoV spikes did not show activity in this assay except for two responders in the HKU1 group.

### Seasonal CoV spike immunity provides negligible protection against SARS-CoV-2 infection

Three weeks after the boost the mice were transduced intranasally with a non-replicating adenovirus expressing human angiotensin converting enzyme 2 (ACE2) followed by a challenge with wild type SARS-CoV-2 strain Washington-1 five days later (15). In this challenge model animals do not suffer severe disease or succumb to infection. However, viral replication in the lung can be assessed to determine protection. A proportion of the animals were euthanized on day 3 or day 5 post challenge and lungs were harvested. The remaining animals were kept to evaluate post-challenge antibody titers. The only group protected from challenge were animals vaccinated with the SARS-CoV-2 spike protein (**Figure 2A and B**). High titers of virus were found in all seasonal CoV spike-vaccinated animals - similar to the negative control group -, with exception of one animal in the OC43 spike group on day 3 and one animal each in the OC43 spike and 229E spike group which displayed lower titers than the control group. None of the animals showed overt disease post challenge. As described, a subset of animals was kept for assessment of the immune response after infection and these animals were bled three weeks post challenge. Anti-SARS-CoV-2 spike titers in all seasonal CoV spike-vaccinated animals had substantially increased (**Figure 2C**). A similar increase was found in the control group with titers in the seasonal CoV spike and the control groups being very similar. Titers in the SARS-CoV-2 spike-immunized animals did not increase post infection suggestive of sterilizing immunity. Little to no ‘back-boosting’ to seasonal CoV spike proteins was detected (**Figure 2D-G**).

**Figure 2:**
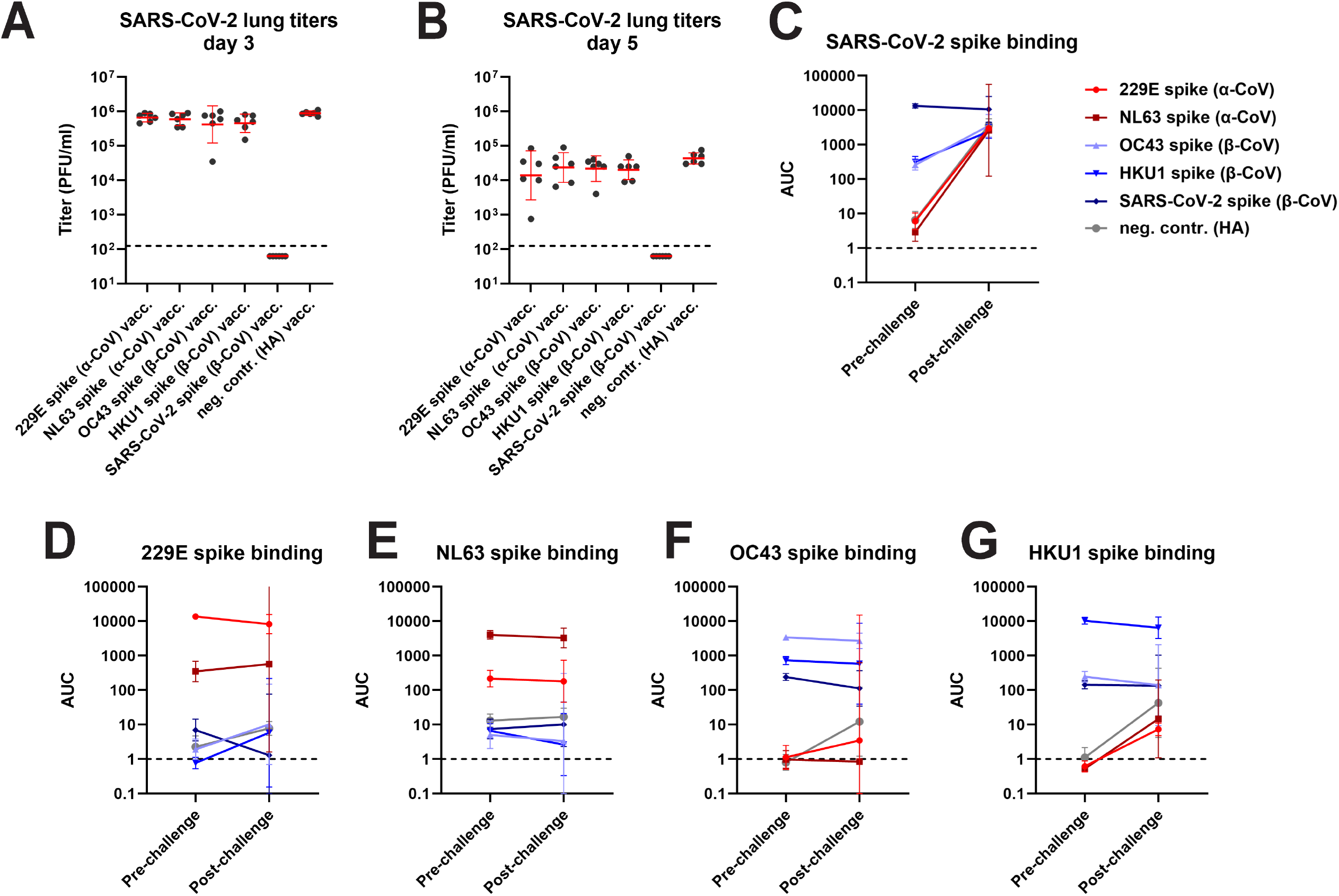
Lung titers of CoV spike vaccinated animals post SARS-CoV-2 challenge and changes in antibody titer. Post challenge lung titers on day 3 and day 5 post challenge are shown in **A** and **B**. Changes in reactivity to SARS-CoV-2 spike, 229E spike, NL63 spike, OC43 spike and HKU1 spike after challenge are shown in **C** to **G**. For A and B, n=6 per group. The n for the pre-challenge time points in C to G are described in the legend of Figure 1. For the post challenge time point, n=3 for all groups except for NL63 where n=2. Significant statistical differences are summarized in Supplemental Table 2 and 3, red bars indicate geometric mean, the error indicates the geometric standard deviation. For the remaining data panels the geometric mean plus 95% confidence intervals are shown. Samples were analyzed once.

### Pre-existing immunity to seasonal CoVs does not negatively impact on protection induced by COVID-19 vaccination

To determine the impact of pre-existing immunity on vaccine-induced protection against SARS-CoV-2, we repeated the vaccination experiment. However, instead of a SARS-CoV-2 challenge, animals were vaccinated 3 weeks post boost with a lipid nanoparticle-formulated nucleoside-modified mRNA vaccine (mRNA-LNP) (16, 17) encoding the SARS-CoV-2 spike protein (**Figure 3A**). An additional control group was added that had not received any vaccination with spike proteins and did not receive the mRNA-LNP vaccine. Vaccination induced high binding and neutralization titers against the SARS-CoV-2 spike in all groups, with the highest titers detected in the SARS-CoV-2 spike protein vaccinated mice and lower titers in the animals vaccinated with seasonal CoV spike or influenza virus HA (**Figure 3B and C**). The similar titers detected in animals pre-immunized with seasonal CoV spikes and influenza virus HA suggest no detrimental ‘original-antigenic sin’-like effect on the immune response to the SARS-CoV-2 mRNA-LNP vaccine. Animals were then transduced with human ACE2 expressing adenovirus and challenged with SARS-CoV-2 as described above. The mice were euthanized on days 3 and 5 post challenge and virus replication in their lungs was assessed. No virus was found in any of the mRNA-LNP-vaccinated animals on either day 3 or 5 (**Figure 3D and E**). However, high virus titers were found in completely naïve animals.

**Figure 3:**
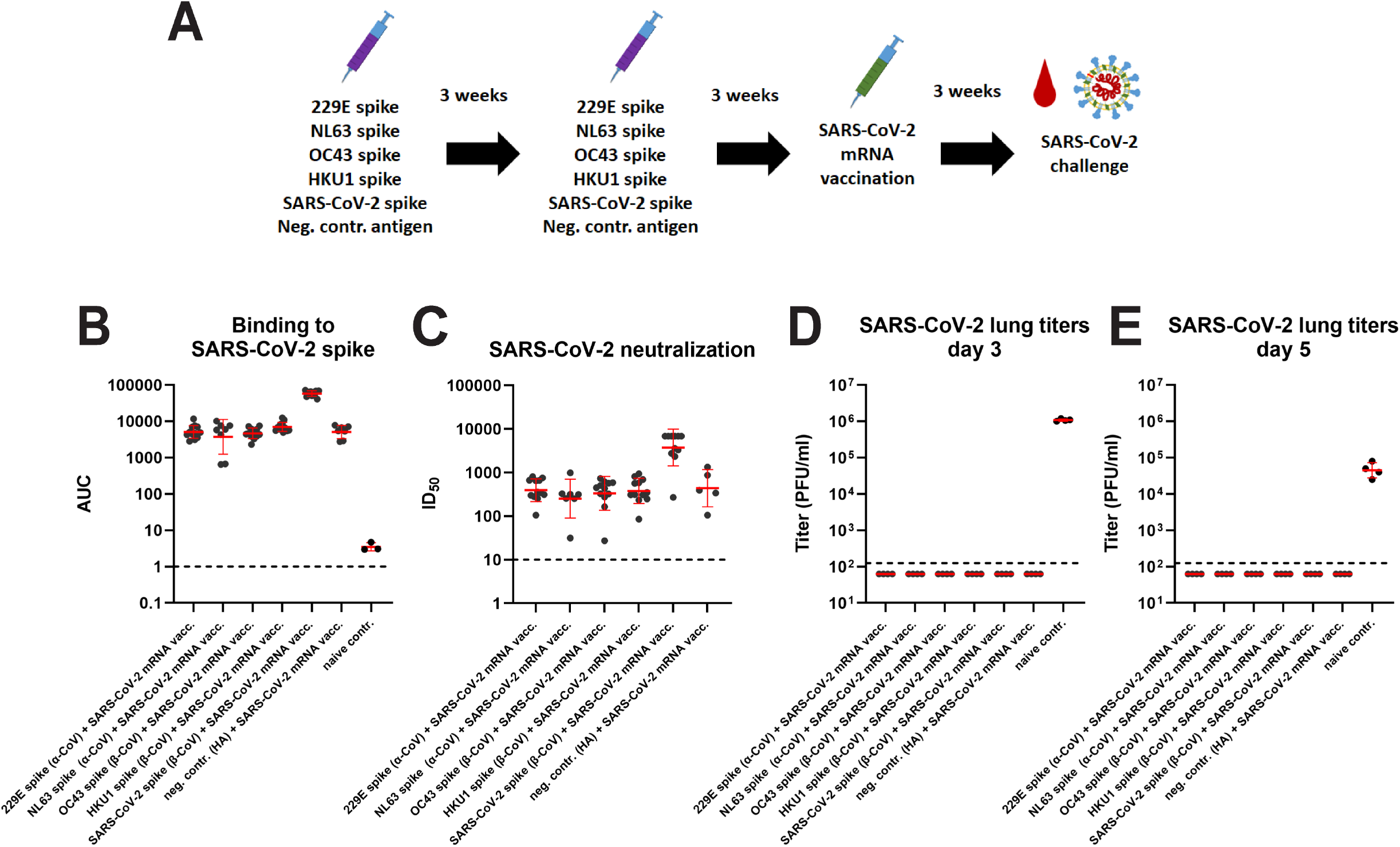
Antibody reactivity post COVID-19 mRNA vaccination and post-challenge lung titers in CoV vaccinated animals. **A** Schematic of vaccination regimen, challenge and sampling. **B** shows binding titers to SARS-CoV-2 spike. **C** shows neutralizing activity to authentic ancestral SARS-CoV-2. Post challenge lung titers on day 3 and day 5 post challenge are shown in **D** and **E**. The n for B was 12 for the 229E, OC43 and HKU1 groups, 8 for the NL63 group, 11 for the SARS-CoV-2 group, 7 for the neg. contr. group and 3 for the naïve group. For C it was 12 for the 229E, OC43 and HKU1 groups, 7 for the NL63 group, 11 for the SARS-CoV-2 group and 5 for the neg. contr. group. For D and E, n=4 per group. Significant statistical differences are summarized in Supplemental Table 4. Red bars indicate geometric mean, the error indicates the geometric standard deviation. Samples were analyzed once.

## Discussion

Pre-existing immunity is known to provide partial protection and can influence immune responses to infection and vaccination for influenza virus (18). A back-boosting or ;original antigenic sin’-like effect on the immune responses to SARS-CoV-2 infection and COVID-19 vaccination has recently been reported as well (13, 14). This was mostly observed towards the β-CoVs OC43 and HKU1 with more reactivity to OC43. Some monoclonal antibodies isolated from exposed individuals, especially the ones targeting the more conserved S2 subunit, have been shown to cross-react to seasonal β-CoV spike proteins as well (19-24). More rare antibodies that target the fusion peptide in S2 even cross-react with α-coronaviruses (25, 26). In addition, it has been hypothesized that antibodies to seasonal CoV spike proteins could, especially in children, provide a degree of protection against SARS-CoV-2 infection and COVID-19 (11).

Here we tested both the hypothesis that pre-existing immunity to seasonal CoV spikes could have a protective effect against SARS-CoV-2 as well as the hypothesis that pre-existing immunity to seasonal CoV spikes could have a detrimental, ‘original antigenic sin’-like impact on protection induced by COVID-19 vaccination. Neither was the case in this mouse model. We did in fact observed a degree of cross-reactivity between β-CoV spikes and between α-CoV spikes as suggested by serological studies and mAb analysis (19-26). This included moderate cross-reactivity of anti-OC43 and anti-HKU1 spike immunity to the SARS-CoV-2 spike. However, this cross-reactivity had no protective effect against SARS-CoV-2 challenge. This suggests that the cross-reactive antibody titers to the spike were either too low to protect in this model or not functional. The lack of ADCC reporter activity despite cross-binding antibodies is in line with this, even though non-neutralizing SARS-CoV-2 spike antibodies are suspected to provide some protection against severe disease with drifted variant viruses in humans. But there may be differences between SARS-CoV-2 specific non-neutralizing antibodies and cross-reactive non-neutralizing antibodies. Furthermore, human and murine antibodies may differ in this respect too. Another interesting observation in this first part of the study was, that challenge with SARS-CoV-2 only increased SARS-CoV-2 spike reactivity but a robust back-boosting effect against β-CoVs as observed in humans (13, 14) was not detected. This is noteworthy since the presence of cross-reactive antibodies in OC43 and HKU1 vaccinated animals was detected pre-challenge and the expectation was that these antibodies would get boosted. While we did not observe any protection conferred by pre-existing immunity against subsequent SARS-CoV-2 challenge, we did also not detected any negative impact on protection induced by a COVID-19 mRNA-LNP vaccine. Mice were fully protected against challenge after mRNA vaccination irrespectively of their pre-existing immunity. While only one challenge dose was used and a challenge dose escalation experiment (had it been possible) could have resulted in differences at more stringent challenge conditions, our data suggest that no strong original antigenic sin-like effect occurs in this model.

Importantly, our study has several limitations. First, the study was performed in mice and this model may just not accurately reflect responses observed in humans, e.g. due to different immunoglobulin germlines and strong differences in length of complementarity determining regions (CDRs) (27). We immunized mice with only one type of spike while humans are exposed to all four seasonal CoVs (7, 8), which could lead to a more cross-reactive immunity. We also did not assess the T-cell response to the different spike proteins and could not evaluate T-cell based immunity to other structural and non-structural proteins of the virus. The seasonal CoVs replicate poorly in cell culture (28, 29) and therefore no neutralization assays were performed.

Despite the lack of a protective effect, it is encouraging that cross-reactivity within the β-CoV spikes could be detected. If this cross-reactivity could be enhanced or if its functionality could be increased in humans, it could lead to broad protection within the β-CoV spikes and could form the basis for a pan-β-CoV vaccine.

## Materials and Methods

### Recombinant proteins

All recombinant proteins were expressed and purified using a mammalian expression system via Expi293F cells (Life Technologies), as described in detail earlier (30, 31). Plasmids to express spike proteins of human coronaviruses which include 229E, HKU1, OC43 and NL63 were generously provided by Dr. Barney Graham. In house expressed proteins were used for immunization. Recombinant proteins were purchased from Sino Biologics (NL63 spike (Cat: 40604-V08B), 2293E spike (Cat: 40605-V08B), HKU1 spike (Cat: 40606-V08B), OC43 spike (Cat: 40607-V08B-B) and SARS-CoV-2 spike (Cat: 40589-V08B1)) to perform ELISAs. Since immunization with in-house expressed proteins led to significant serum reactivity against the trimerization domain, proteins from Sino Biologicals (with no trimerization domain) were purchased and used. Hence, the immune response to the respective proteins could be assessed in a more accurate fashion.

### mRNA-LNP vaccine production

The mRNA vaccine was designed and made as described (16, 17). Briefly, the coding sequence of full length Δfurin spike protein (RRAR furin cleavage site abolished between amino acids 682-685) from the Wuhan-1 strain (Wuhan-Hu-1, GenBank: MN908947.3) was codon-optimized, synthesized and cloned into an mRNA production plasmid. The spike-encoding mRNA was transcribed to contain 101 nucleotide-long poly(A) tails. m1Ψ-5’-triphosphate (TriLink) instead of UTP was used to generate modified nucleoside-containing mRNA. Capping of the *in vitro* transcribed mRNAs was performed co-transcriptionally using the trinucleotide cap1 analog, CleanCap (TriLink). mRNA was purified by cellulose purification as described (32). The mRNA was analyzed by agarose gel electrophoresis and was stored frozen at -20°C. The cellulose-purified m1Ψ-containing mRNA was encapsulated in LNPs using a self-assembly process as previously described wherein an ethanolic lipid mixture of ionizable cationic lipid, phosphatidylcholine, cholesterol and polyethylene glycol-lipid was rapidly mixed with an aqueous solution containing mRNA at acidic pH. The RNA-loaded particles were characterized and subsequently stored at -80°C at a concentration of 1 μ (33)g μl^-1^. The mean hydrodynamic diameter of these mRNA-LNP was ∼80 nm with a polydispersity index of 0.02-0.06 and an encapsulation efficiency of ∼95%.

### Cells and viruses

Vero.E6 cells (ATCC CRL-1586, clone E6) were maintained for cell culture using Dulbecco’s modified Eagle medium (Gibco), which was supplemented with Antibiotic-Antimycotic (100 U/ml penicillin–100 μg/ml streptomycin–0.25 μg/ml amphotericin B; Gibco), 10% of fetal bovine serum (FBS; Corning), and 1% HEPES (N-2-hydroxyethylpiperazine-N-2-ethane sulfonic acid; Gibco). Wild type SARS-CoV-2 (isolate USA-WA1/2020) was grown in Vero.E6 cells for 3 days at 37°C and then the supernatant was clarified via centrifugation at 1,000g for 10 minutes. Virus stocks were stored at −80°C. The protocol is described in greater detail previously (34, 35). All work with authentic live SARS-CoV-2 was performed in the biosafety level 3 (BSL-3) facility following institutional guidelines.

### *In vivo* studies

All animal procedures were performed by following the Institutional Animal Care and Use Committee (IACUC) guidelines of the Icahn School of Medicine at Mount Sinai IACUC and according to an approved protocol. Six- to 8-week-old female, BALB/c mice were vaccinated via the intramuscular route with 3 μg of each respective protein with 1:1 mixture of Addavax (Invivogen) in a total volume of 50 μL. After 3 weeks, mice were bled and vaccinated with the same protein as original the vaccination. After 3 weeks, mice were administered anesthesia via the intraperitoneal route and intranasally transduced with AdV-hACE2 at 2.5 × 10^8^ plaque-forming units (PFUs) per mouse (15). Anesthesia was prepared using 0.15 mg/kg of body weight ketamine and 0.03 mg/kg xylazine in water. Five days later, all mice were infected with authentic wild-type SARS-CoV-2 intranasally with 1 × 10^5^ PFU. Mice were humanely euthanized on day 3 and day 5 for assessment of virus in the lungs. Lungs were homogenized using a BeadBlaster 24 (Benchmark) homogenizer (36-38). Viral load in the lung was quantified via a classic plaque assay (39, 40). For mice that also received mRNA-LNP vaccine, the vaccine was administered via the intramuscular route and each mouse was immunized with a single dose of 3 ug of mRNA-LNP.

### ELISA

Ninety-six-well plates (Immulon 4 HBX; Thermo Fisher Scientific) were coated with 2 μg/mL of each antigen with 50 μL/well at 4°C overnight for 16 hours. The following day, the coating solution was removed, and each plate was blocked with 100 μL/well of 3% non-fat milk (American Bio; catalog no. AB10109-01000) in phosphate buffered saline containing 0.01% Tween 20 (PBS-T). Blocking solution was incubated on the plates for 1 hour at room temperature (RT). Serum samples were tested starting at a dilution of 1:50 with 1:5-fold subsequent serial dilutions. Serum samples were added to the plates for 2 hours at RT. Next, the plates were properly washed 3 times with 200 μL/well of PBS-T. Anti-mouse IgG-horseradish peroxidase (HRP)-conjugated antibody (Rockland; catalog no. 610–4302) was used at a dilution of 1:3,000 in 1% non-fat milk in PBS-T, and 100 μL of the prepared solution was added on to each well for 1 hour at RT. The plates were washed three times with 200 μL/well of PBS-T and patted on paper towels to dry. Developing solution was made in sterile water (WFI; Gibco) using SigmaFast OPD (o-phenylenediamine dihydrochloride, catalog no. P9187; Sigma-Aldrich), and 100 μL was added to each well for a total of 10 minutes. To stop the reaction, 50 μL/well of 3 M hydrochloric acid was added, and the plates were read in a Synergy 4 (BioTek) plate reader at an absorbance of 490 nanometers. Data were analyzed in GraphPad Prism 8. An area under the curve was calculate for the dilution series and reported. This protocol is described in earlier literature in greater detail (34, 41-43).

### Neutralization assays

Vero.E6 cells were seeded in a 96-well cell culture plate (Corning; 3340) 1 day prior to the assay, at a density of 20,000 cells per well. Mouse serum samples were heat inactivated at 56°C for 1 hour prior to use. Serum dilutions were prepared in 1× minimal essential medium (MEM; Gibco) supplemented with 1% FBS. Virus was diluted to 10,000 50% tissue culture infectious doses (TCID_50_s)/mL, and 80 μL of virus and 80 μL of serum were incubated together for 1 hour at RT. After the incubation, 120 μL of virus–serum mixture was used to infect cells for 1 hour at 37°C. Next, the virus– serum mix was removed and 100 μL of each corresponding dilution was added to each well. A volume of 100 μL of 1X MEM were also added to the plates to get to a total volume of 200 μL in each well. Cells were incubated at 37°C for 3 days and later fixed with 10% paraformaldehyde (Polysciences) for 24 hours. After 24 hours, the paraformaldehyde solution was discarded and cells were permeabilized for intracellular staining. Staining and quantification were performed as previously described (30, 34, 44, 45).

### ADCC reporter assay

An assay similar to a setup previously published for influenza virus was set up to measure ADCC reporter activity (46). Vero.E6 cells were seeded in white 96-well cultures plates (Corning; 3917) at a density of 20,000 cells per well and infected 24 hours later with recombinant Newcastle disease virus (NDV) expressing the prefusion-stabilized spike protein from SARS-CoV-2 (NDV-HXP-S) (47, 48) at a multiplicity of infection (MOI) of 2. After 1 hour of incubation the infection inoculum was removed and replaced with 100 μl Roswell Park Memorial Institute (RPMI) 1640 media (Cytiva; SH30027.02) supplemented with 2% super low-IgG FBS (Cytiva; SH30898.02HI). Heat inactivated mouse serum samples were serially diluted 1:3 in RPMI 1640 media supplemented with 2% super low-IgG FBS from a starting dilution of 1:10. Forty-eight hours post-infection, the media was removed and 25ul RPMI 1640 supplemented with 2% super low-IgG FBS was added in addition to 25ul of diluted sera. ADCC bioassay effector cells (Promega; M1201) expressing the murine FcγRIV receptor were added to each well at a density of 75,000 cells per well in a volume of 25 μl. The effector cells and serum were incubated with the cells for 6 hours after which 75ul Bio-Glo reagent (Promega; G7940) was added to each well. Plates were incubated at RT for 10 minutes in the dark before luminescence was measured using a Synergy 4 (BioTek) plate reader. An area under the curve was calculate for the dilution series and reported.

### Statistical analysis

Groups were compared using a one-sided ANOVA corrected for multiple comparisons after log transformation in. Pre- and post-challenge titers were compared using a t-test after log transformation. Analysis was performed in GraphPad Prism (version 9.0.1). Results from statistical testing are reported in Supplementary Tables 1-4.

## Acknowledgments

We would like to thank Dr. Randy A. Albrecht for oversight of the conventional BSL3 biocontainment facility, which makes our work with live SARS-CoV-2 possible. This work was partially funded by the Centers of Excellence for Influenza Research and Surveillance (CEIRS, contract # HHSN272201400008C), by the Collaborative Influenza Vaccine Innovation Centers (CIVICs contract # 75N93019C00051) and by the generous support by anonymous donors. LNP formulation of mRNAs was performed by Acuitas Therapeutics, Vancouver, BC Canada.

## Data availability statement

Data will be made publicly available upon publication and upon request for peer review.

## Supplemental Tables

**Supplemental Table 1.**

| Group comparison using Tukey's multiple comparisons test | P value |
| --- | --- |
| Figure 1B |  |
| 229E spike (α-CoV) vacc. vs. NL63 spike (α-CoV) vacc. | <0.0001 |
| 229E spike (α-CoV) vacc. vs. OC43 spike (β-CoV) vacc. | <0.0001 |
| 229E spike (α-CoV) vacc. vs. HKU1 spike (β-CoV) vacc. | <0.0001 |
| 229E spike (α-CoV) vacc. vs. SARS-CoV-2 spike (β-CoV) vacc. | <0.0001 |
| 229E spike (α-CoV) vacc. vs. neg. contr. (HA) vacc. | <0.0001 |
| NL63 spike (α-CoV) vacc. vs. OC43 spike (β-CoV) vacc. | <0.0001 |
| NL63 spike (α-CoV) vacc. vs. HKU1 spike (β-CoV) vacc. | <0.0001 |
| NL63 spike (α-CoV) vacc. vs. SARS-CoV-2 spike (β-CoV) vacc. | <0.0001 |
| NL63 spike (α-CoV) vacc. vs. neg. contr. (HA) vacc. | <0.0001 |
| OC43 spike (β-CoV) vacc. vs. HKU1 spike (β-CoV) vacc. | 0.2214 |
| OC43 spike (β-CoV) vacc. vs. SARS-CoV-2 spike (β-CoV) vacc. | 0.0268 |
| OC43 spike (β-CoV) vacc. vs. neg. contr. (HA) vacc. | 0.9984 |
| HKU1 spike (β-CoV) vacc. vs. SARS-CoV-2 spike (β-CoV) vacc. | <0.0001 |
| HKU1 spike (β-CoV) vacc. vs. neg. contr. (HA) vacc. | 0.0772 |
| SARS-CoV-2 spike (β-CoV) vacc. vs. neg. contr. (HA) vacc. | 0.0747 |
| Figure 1C |  |
| 229E spike (α-CoV) vacc. vs. NL63 spike (α-CoV) vacc. | <0.0001 |
| 229E spike (α-CoV) vacc. vs. OC43 spike (β-CoV) vacc. | <0.0001 |
| 229E spike (α-CoV) vacc. vs. HKU1 spike (β-CoV) vacc. | <0.0001 |
| 229E spike (α-CoV) vacc. vs. SARS-CoV-2 spike (β-CoV) vacc. | <0.0001 |
| 229E spike (α-CoV) vacc. vs. neg. contr. (HA) vacc. | <0.0001 |
| NL63 spike (α-CoV) vacc. vs. OC43 spike (β-CoV) vacc. | <0.0001 |
| NL63 spike (α-CoV) vacc. vs. HKU1 spike (β-CoV) vacc. | <0.0001 |
| NL63 spike (α-CoV) vacc. vs. SARS-CoV-2 spike (β-CoV) vacc. | <0.0001 |
| NL63 spike (α-CoV) vacc. vs. neg. contr. (HA) vacc. | <0.0001 |
| OC43 spike (β-CoV) vacc. vs. HKU1 spike (β-CoV) vacc. | 0.9820 |
| OC43 spike (β-CoV) vacc. vs. SARS-CoV-2 spike (β-CoV) vacc. | 0.9179 |
| OC43 spike (β-CoV) vacc. vs. neg. contr. (HA) vacc. | 0.1729 |
| HKU1 spike (β-CoV) vacc. vs. SARS-CoV-2 spike (β-CoV) vacc. | 0.9995 |
| HKU1 spike (β-CoV) vacc. vs. neg. contr. (HA) vacc. | 0.4668 |
| SARS-CoV-2 spike (β-CoV) vacc. vs. neg. contr. (HA) vacc. | 0.6841 |
| Figure 1D |  |
| 229E spike (α-CoV) vacc. vs. NL63 spike (α-CoV) vacc. | 0.9981 |
| 229E spike (α-CoV) vacc. vs. OC43 spike (β-CoV) vacc. | <0.0001 |
| 229E spike (α-CoV) vacc. vs. HKU1 spike (β-CoV) vacc. | <0.0001 |
| 229E spike (α-CoV) vacc. vs. SARS-CoV-2 spike (β-CoV) vacc. | <0.0001 |
| 229E spike (α-CoV) vacc. vs. neg. contr. (HA) vacc. | 0.9180 |
| NL63 spike (α-CoV) vacc. vs. OC43 spike (β-CoV) vacc. | <0.0001 |
| NL63 spike (α-CoV) vacc. vs. HKU1 spike (β-CoV) vacc. | <0.0001 |
| NL63 spike (α-CoV) vacc. vs. SARS-CoV-2 spike (β-CoV) vacc. | <0.0001 |
| NL63 spike (α-CoV) vacc. vs. neg. contr. (HA) vacc. | 0.9908 |
| OC43 spike (β-CoV) vacc. vs. HKU1 spike (β-CoV) vacc. | 0.0002 |
| OC43 spike (β-CoV) vacc. vs. SARS-CoV-2 spike (β-CoV) vacc. | <0.0001 |
| OC43 spike (β-CoV) vacc. vs. neg. contr. (HA) vacc. | <0.0001 |
| HKU1 spike (β-CoV) vacc. vs. SARS-CoV-2 spike (β-CoV) vacc. | 0.0090 |
| HKU1 spike (β-CoV) vacc. vs. neg. contr. (HA) vacc. | <0.0001 |
| SARS-CoV-2 spike (β-CoV) vacc. vs. neg. contr. (HA) vacc. | <0.0001 |
| Figure 1E |  |
| 229E spike (α-CoV) vacc. vs. NL63 spike (α-CoV) vacc. | 0.9809 |
| 229E spike (α-CoV) vacc. vs. OC43 spike (β-CoV) vacc. | <0.0001 |
| 229E spike (α-CoV) vacc. vs. HKU1 spike (β-CoV) vacc. | <0.0001 |
| 229E spike (α-CoV) vacc. vs. SARS-CoV-2 spike (β-CoV) vacc. | <0.0001 |
| 229E spike (α-CoV) vacc. vs. neg. contr. (HA) vacc. | 0.1351 |
| NL63 spike (α-CoV) vacc. vs. OC43 spike (β-CoV) vacc. | <0.0001 |
| NL63 spike (α-CoV) vacc. vs. HKU1 spike (β-CoV) vacc. | <0.0001 |
| NL63 spike (α-CoV) vacc. vs. SARS-CoV-2 spike (β-CoV) vacc. | <0.0001 |
| NL63 spike (α-CoV) vacc. vs. neg. contr. (HA) vacc. | 0.0187 |
| OC43 spike (β-CoV) vacc. vs. HKU1 spike (β-CoV) vacc. | <0.0001 |
| OC43 spike (β-CoV) vacc. vs. SARS-CoV-2 spike (β-CoV) vacc. | 0.2235 |
| OC43 spike (β-CoV) vacc. vs. neg. contr. (HA) vacc. | <0.0001 |
| HKU1 spike (β-CoV) vacc. vs. SARS-CoV-2 spike (β-CoV) vacc. | <0.0001 |
| HKU1 spike (β-CoV) vacc. vs. neg. contr. (HA) vacc. | <0.0001 |
| SARS-CoV-2 spike (β-CoV) vacc. vs. neg. contr. (HA) vacc. | <0.0001 |
| Figure 1F |  |
| 229E spike (α-CoV) vacc. vs. NL63 spike (α-CoV) vacc. | 0.1735 |
| 229E spike (α-CoV) vacc. vs. OC43 spike (β-CoV) vacc. | <0.0001 |
| 229E spike (α-CoV) vacc. vs. HKU1 spike (β-CoV) vacc. | <0.0001 |
| 229E spike (α-CoV) vacc. vs. SARS-CoV-2 spike (β-CoV) vacc. | <0.0001 |
| 229E spike (α-CoV) vacc. vs. neg. contr. (HA) vacc. | 0.9998 |
| NL63 spike (α-CoV) vacc. vs. OC43 spike (β-CoV) vacc. | <0.0001 |
| NL63 spike (α-CoV) vacc. vs. HKU1 spike (β-CoV) vacc. | <0.0001 |
| NL63 spike (α-CoV) vacc. vs. SARS-CoV-2 spike (β-CoV) vacc. | <0.0001 |
| NL63 spike (α-CoV) vacc. vs. neg. contr. (HA) vacc. | 0.1018 |
| OC43 spike (β-CoV) vacc. vs. HKU1 spike (β-CoV) vacc. | 0.9861 |
| OC43 spike (β-CoV) vacc. vs. SARS-CoV-2 spike (β-CoV) vacc. | <0.0001 |
| OC43 spike (β-CoV) vacc. vs. neg. contr. (HA) vacc. | <0.0001 |
| HKU1 spike (β-CoV) vacc. vs. SARS-CoV-2 spike (β-CoV) vacc. | <0.0001 |
| HKU1 spike (β-CoV) vacc. vs. neg. contr. (HA) vacc. | <0.0001 |
| SARS-CoV-2 spike (β-CoV) vacc. vs. neg. contr. (HA) vacc. | <0.0001 |
| Figure 1G |  |
| 229E spike (α-CoV) vacc. vs. NL63 spike (α-CoV) vacc. | >0.9999 |
| 229E spike (α-CoV) vacc. vs. OC43 spike (β-CoV) vacc. | >0.9999 |
| 229E spike (α-CoV) vacc. vs. HKU1 spike (β-CoV) vacc. | 0.6268 |
| 229E spike (α-CoV) vacc. vs. SARS-CoV-2 spike (β-CoV) vacc. | <0.0001 |
| 229E spike (α-CoV) vacc. vs. neg. contr. (HA) vacc. | >0.9999 |
| NL63 spike (α-CoV) vacc. vs. OC43 spike (β-CoV) vacc. | >0.9999 |
| NL63 spike (α-CoV) vacc. vs. HKU1 spike (β-CoV) vacc. | 0.6268 |
| NL63 spike (α-CoV) vacc. vs. SARS-CoV-2 spike (β-CoV) vacc. | <0.0001 |
| NL63 spike (α-CoV) vacc. vs. neg. contr. (HA) vacc. | >0.9999 |
| OC43 spike (β-CoV) vacc. vs. HKU1 spike (β-CoV) vacc. | 0.6268 |
| OC43 spike (β-CoV) vacc. vs. SARS-CoV-2 spike (β-CoV) vacc. | <0.0001 |
| OC43 spike (β-CoV) vacc. vs. neg. contr. (HA) vacc. | >0.9999 |
| HKU1 spike (β-CoV) vacc. vs. SARS-CoV-2 spike (β-CoV) vacc. | 0.0063 |
| HKU1 spike (β-CoV) vacc. vs. neg. contr. (HA) vacc. | 0.6268 |
| SARS-CoV-2 spike (β-CoV) vacc. vs. neg. contr. (HA) vacc. | <0.0001 |

**Supplemental Table 2.**

| Group comparison using Tukey's multiple comparisons test | P value |
| --- | --- |
| Figure 2A |  |
| 229E spike (α-CoV) vacc. vs. NL63 spike (α-CoV) vacc. | 0.9994 |
| 229E spike (α-CoV) vacc. vs. OC43 spike (β-CoV) vacc. | 0.7767 |
| 229E spike (α-CoV) vacc. vs. HKU1 spike (β-CoV) vacc. | 0.8858 |
| 229E spike (α-CoV) vacc. vs. SARS-CoV-2 spike (β-CoV) vacc. | <0.0001 |
| 229E spike (α-CoV) vacc. vs. neg. contr. (HA) vacc. | 0.9606 |
| NL63 spike (α-CoV) vacc. vs. OC43 spike (β-CoV) vacc. | 0.9194 |
| NL63 spike (α-CoV) vacc. vs. HKU1 spike (β-CoV) vacc. | 0.9733 |
| NL63 spike (α-CoV) vacc. vs. SARS-CoV-2 spike (β-CoV) vacc. | <0.0001 |
| NL63 spike (α-CoV) vacc. vs. neg. contr. (HA) vacc. | 0.8557 |
| OC43 spike (β-CoV) vacc. vs. HKU1 spike (β-CoV) vacc. | 0.9999 |
| OC43 spike (β-CoV) vacc. vs. SARS-CoV-2 spike (β-CoV) vacc. | <0.0001 |
| OC43 spike (β-CoV) vacc. vs. neg. contr. (HA) vacc. | 0.2971 |
| HKU1 spike (β-CoV) vacc. vs. SARS-CoV-2 spike (β-CoV) vacc. | <0.0001 |
| HKU1 spike (β-CoV) vacc. vs. neg. contr. (HA) vacc. | 0.4196 |
| SARS-CoV-2 spike (β-CoV) vacc. vs. neg. contr. (HA) vacc. | <0.0001 |
| Figure 2C |  |
| 229E spike (α-CoV) vacc. vs. NL63 spike (α-CoV) vacc. | 0.9171 |
| 229E spike (α-CoV) vacc. vs. OC43 spike (β-CoV) vacc. | 0.9569 |
| 229E spike (α-CoV) vacc. vs. HKU1 spike (β-CoV) vacc. | 0.9806 |
| 229E spike (α-CoV) vacc. vs. SARS-CoV-2 spike (β-CoV) vacc. | <0.0001 |
| 229E spike (α-CoV) vacc. vs. neg. contr. (HA) vacc. | 0.2900 |
| NL63 spike (α-CoV) vacc. vs. OC43 spike (β-CoV) vacc. | >0.9999 |
| NL63 spike (α-CoV) vacc. vs. HKU1 spike (β-CoV) vacc. | 0.9997 |
| NL63 spike (α-CoV) vacc. vs. SARS-CoV-2 spike (β-CoV) vacc. | <0.0001 |
| NL63 spike (α-CoV) vacc. vs. neg. contr. (HA) vacc. | 0.8522 |
| OC43 spike (β-CoV) vacc. vs. HKU1 spike (β-CoV) vacc. | >0.9999 |
| OC43 spike (β-CoV) vacc. vs. SARS-CoV-2 spike (β-CoV) vacc. | <0.0001 |
| OC43 spike (β-CoV) vacc. vs. neg. contr. (HA) vacc. | 0.7784 |
| HKU1 spike (β-CoV) vacc. vs. SARS-CoV-2 spike (β-CoV) vacc. | <0.0001 |
| HKU1 spike (β-CoV) vacc. vs. neg. contr. (HA) vacc. | 0.6963 |
| SARS-CoV-2 spike (β-CoV) vacc. vs. neg. contr. (HA) vacc. | <0.0001 |

**Supplemental Table 3.**

| Pre/post challenge comparison, t-test | P value |
| --- | --- |
| Figure 2C |  |
| 229E spike (α-CoV) | <0.0001 |
| NL63 spike (α-CoV) | <0.0001 |
| OC43 spike (β-CoV) | <0.0001 |
| HKU1 spike (β-CoV) | 0.0003 |
| SARS-CoV-2 spike (β-CoV) | 0.2010 |
| neg. contr. (HA) | <0.0001 |
| Figure 2D |  |
| 229E spike (α-CoV) | 0.0046 |
| NL63 spike (α-CoV) | 0.6519 |
| OC43 spike (β-CoV) | 0.0507 |
| HKU1 spike (β-CoV) | 0.0014 |
| SARS-CoV-2 spike (β-CoV) | 0.0917 |
| neg. contr. (HA) | 0.1586 |
| Figure 2E |  |
| 229E spike (α-CoV) | 0.7836 |
| NL63 spike (α-CoV) | 0.6472 |
| OC43 spike (β-CoV) | 0.7076 |
| HKU1 spike (β-CoV) | 0.1493 |
| SARS-CoV-2 spike (β-CoV) | 0.7036 |
| neg. contr. (HA) | 0.6119 |
| Figure 2F |  |
| 229E spike (α-CoV) | 0.3442 |
| NL63 spike (α-CoV) | 0.8733 |
| OC43 spike (β-CoV) | 0.1553 |
| HKU1 spike (β-CoV) | 0.5709 |
| SARS-CoV-2 spike (β-CoV) | 0.0212 |
| neg. contr. (HA) | 0.0004 |
| Figure 2G |  |
| 229E spike (α-CoV) | <0.0001 |
| NL63 spike (α-CoV) | <0.0001 |
| OC43 spike (β-CoV) | 0.2383 |
| HKU1 spike (β-CoV) | 0.1213 |
| SARS-CoV-2 spike (β-CoV) | 0.8895 |
| neg. contr. (HA) | 0.0002 |

**Supplemental Table 4.**
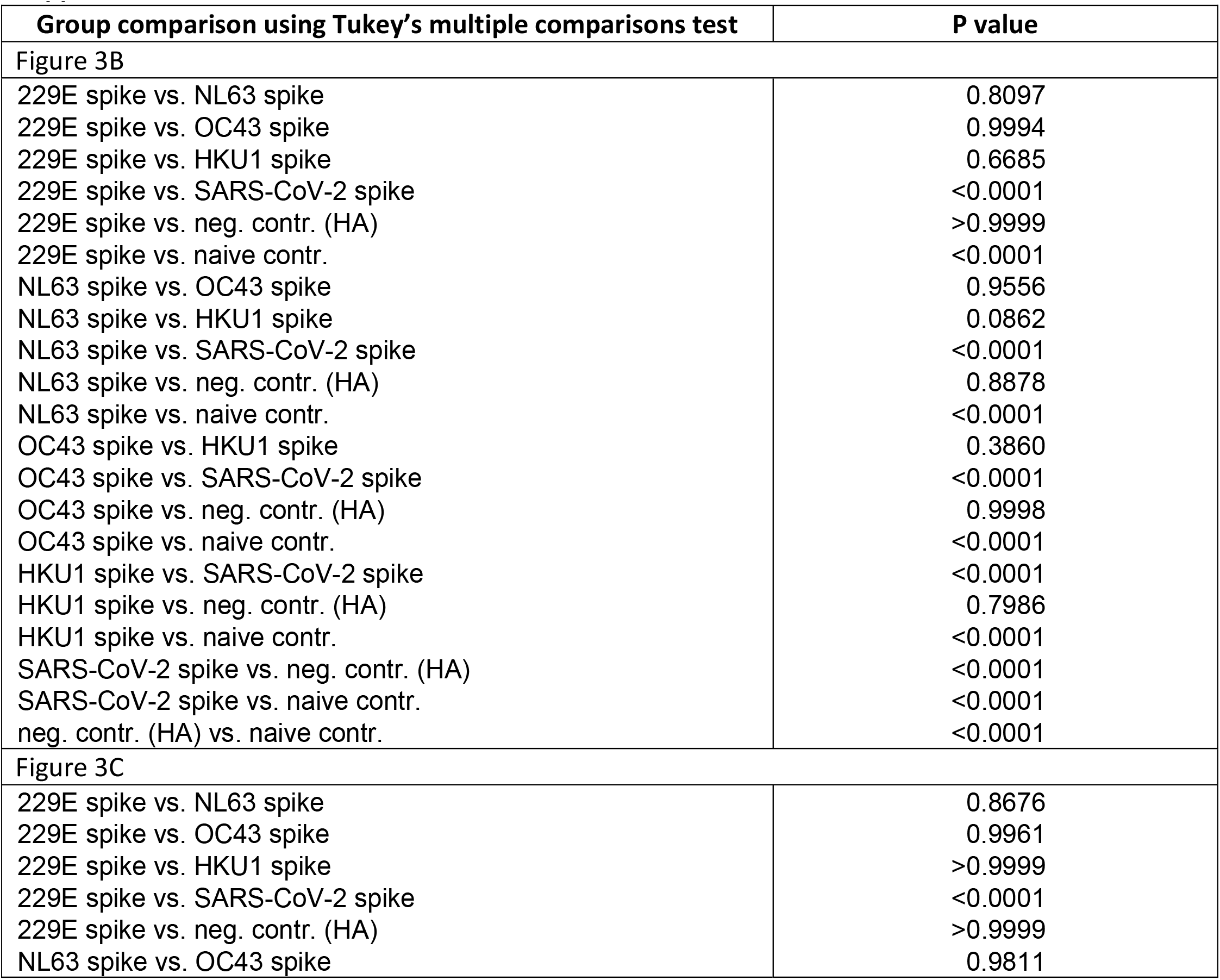
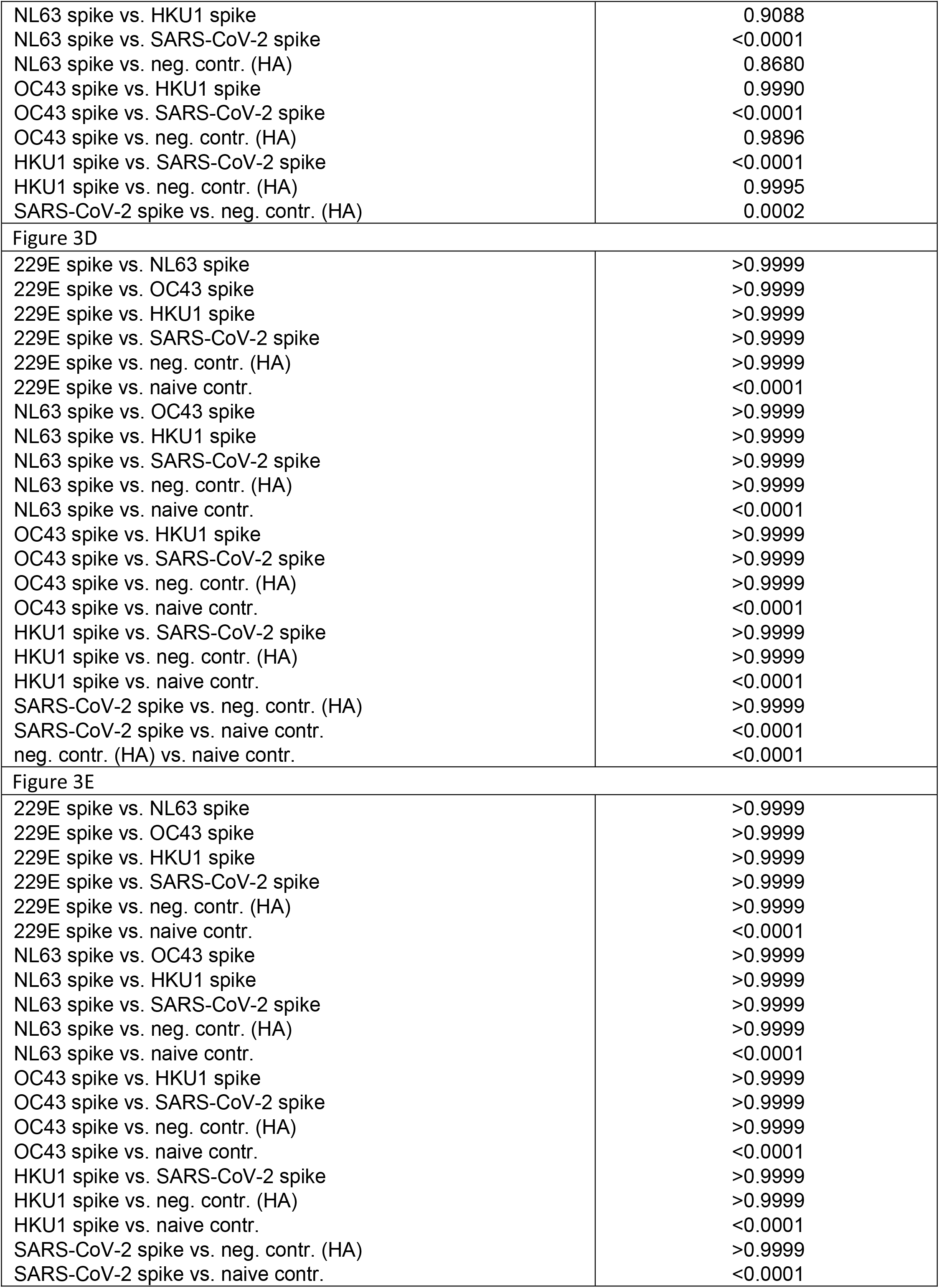
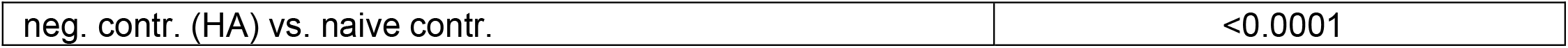

